# Does the stress axis mediate behavioural flexibility in a social cichlid, *Neolamprologus pulcher*?

**DOI:** 10.1101/2023.09.07.556642

**Authors:** Stefan Fischer, Zala Ferlinc, Katharina Hirschenhauser, Barbara Taborsky, Leonida Fusani, Sabine Tebbich

**Affiliations:** Konrad Lorenz Institute for Ethology, Department of Interdisciplinary Life Sciences, University of Veterinary Medicine Vienna, Savoyenstrasse 1, 1160 Vienna, Austria; Department of Behavioral and Cognitive Biology, University of Vienna, Djerassiplatz 1, 1030 Vienna, Austria; University College for Education of Upper Austria (PH OÖ), Kaplanhofstraße 40, 4020 Linz, Austria; Division of Behavioural Ecology, University of Bern, Wohlenstrasse 50a, CH-3032 Hinterkappelen, Switzerland

**Keywords:** cortisol, glucocorticoid receptor, mifepristone, RU 484, Detour task, predator, *Lepidiolamprologus elongatus*, learning

## Abstract

Behavioural flexibility plays a major role in the way animals cope with novel situations, and physiological stress responses are adaptive and highly efficient mechanisms to cope with unpredictable events. Previous studies investigating the role of stress responses in mediating behavioural flexibility were mostly done in laboratory rodents using stressors and cognitive challenges unrelated to the ecology of the species. To better understand how stress mediates behavioural flexibility in a natural context, direct manipulations of the stress response and cognitive tests in ecologically relevant contexts are needed. To this aim, we pharmacologically blocked glucocorticoid receptors (GR) in adult *Neolamprologus pulcher* using a minimally invasive application of a GR antagonist. GR blockade prevents the recovery after a stressful event, which we predicted to impair behavioural flexibility. After the application of the GR antagonist, we repeatedly exposed fish to a predator and tested their behavioural flexibility using a detour task, i.e. fish had to find a new, longer route to the shelter when the shortest route was blocked. While the latencies to find the shelter were not different between treatments, GR blocked fish showed more failed attempts during the detour tasks than control fish. Furthermore, weak performance during the detour tasks was accompanied by an increase of fear related behaviours. This suggests that blocking GR changed the perception of fear and resulted in an impaired behavioural flexibility. Therefore, our results support a causal link between the capacity to recover from stressors and behavioural flexibility in *N. pulcher* with potential consequences for an effective and adaptive coping with changing environments.

## Introduction

Stress is a ubiquitous feature of life and all organisms have evolved internal mechanisms to cope with external stressors (Taborsky et al., 2021). Stressors can be abiotic or biotic, ranging from extreme temperatures to predator attacks or social stressors such as aggressive conflicts. An adequate response to a stressor, the so-called stress response, has major fitness consequences because it is of utmost importance for the survival of the individual (Lemonnier et al., 2022; McEwen and Wingfield, 2003; Romero et al., 2009; Sopinka et al., 2015).

When a stressor is experienced the adaptive stress response in vertebrates is generally characterised by a physiological increase of circulating glucocorticoids (GCs) (Sapolsky et al., 2000). Cortisol is the major GC and a main actor of the stress response in fish and most mammals (Sopinka et al., 2015). Cortisol can cross the blood-brain barrier where it binds to high affinity mineralcorticoid receptors (MRs) and lower affinity glucocorticoid receptors (GRs) that are expressed across the brain (Joels et al., 2008; Roozendaal et al., 2009). Cortisol acts on GRs and MRs to induce a negative feedback that reduces the further release of cortisol, which slowly returns to baseline levels (de Kloet, 2014; Joels et al., 2008). In contrast to mammals, most teleosts have two GRs with different hormone sensitivities (Bury, 2017). MRs also play a different role in teleosts because MR mediates the effects of aldosterone, which is absent in fish (Faught and Vijayan, 2018). The swift increase and slow decrease of cortisol is an adaptive stress response leading to beneficial changes in physiology, behaviour and cognition to successfully cope with the stressor. On the contrary, if cortisol concentrations do not return to baseline levels after the stressor ends, prolonged elevation of cortisol has adverse effects on health and behaviour particularly on cognitive abilities (Sandi and Pinelo-Nava, 2007; Schwabe et al., 2012).

The flexible adjustment of behaviours, or behavioural flexibility, requires different cognitive abilities including learning and the seeking of alternative solutions (Sol and Lefebvre, 2000; Tebbich et al., 2010). To find an alternative solution animals have to control and supress a predisposition for a behavioural response to show a more appropriate behaviour (i.e. inhibitory control), which is a core executive function and is beneficial to animals in changing environments (Burkart et al., 2017; Diamond, 2013; Triki et al., 2023). GR activity has been shown to play an important role in influencing cognitive function and particularly in regulating learning performances (for reviews see: de Kloet et al., 2005; Roozendaal et al., 2009; Sandi and Pinelo-Nava, 2007). Previous studies using laboratory rodents have shown that GRs play a causal role in determining memory formation (Oitzl and de Kloet, 1992; Roozendaal and McGaugh, 1997), fear conditioning (Donley et al., 2005), and reversal learning, a commonly used proxy for behavioural flexibility (Bryce and Howland, 2015). Studies that tested the effects of the stress axis on behavioural flexibility in adult individuals of various species manipulated mostly type or duration of stressor (e.g. Bryce and Howland, 2015; Butts et al., 2013; Calandreau et al., 2011; George et al., 2015; Graybeal et al., 2014; Graybeal et al., 2011; Thai et al., 2013). Some studies have investigated the role of GR activitiy for learning and behavioural flexibility by manipulating GR mediated action. For example, laboratory rats receiving the specific GR antagonist mifepristone performed worse in a spatial learning task than rats receiving a control treatment (Oitzl and de Kloet, 1992). Other studies have focused on functional consequences of experiencing stressors and the resulting variation in cognition without assessing the underlying physiological mechanisms (Ferrari, 2014; Potvin, 2017; Pouca et al., 2021). To reveal that adaptive stress responses help animals to cope with changing environments we need to directly link physiological parameters involved in the recovery from stressors with the flexible adjustment of behaviour.

Fear conditioning and fear related behaviours are tightly linked with the activity of GR as well as GC concentrations (Schulkin et al., 2005a). Particularly GRs play an important role in the retention of fear memory and fear extinction after repeated exposures to a stressor (Cordero and Sandi, 1998; Kolber et al., 2008; Korte, 2001). For example, chronically stressed rats receiving a GC antagonist displayed more fear related behaviour to the conditioned stimulus alone compared to chronically stressed rats receiving a vehicle injection (Conrad et al., 2004).

In this work, we administered with a minimally invasive method a high dose of a GR antagonist (mifepristone) to adult *Neolamprologus pulcher* and tested their behavioural flexibility in a spatial task after being repeatedly exposed to a predator. Mifepristone is an antiprogestin and GR antagonist with several clinical applications (Spitz, 2003). At higher dosages it mainly acts as a GR antagonist (Alderman et al., 2012; Carbajal et al., 2023; Lawrence et al., 2017; Molitch, 2022; Oitzl and de Kloet, 1992; Ros et al., 2012; Veillette et al., 2007) with an affinity to GRs that is tenfold higher than that of cortisol (Molitch, 2022). In contrast, at lower dosages it mainly acts as an antiprogestin interacting with progesterone receptors (PRs) and as such, it is also used as a non-invasive method to terminate pregnancies (Blüthgen et al., 2013; Spitz et al., 1996). In the spatial task, individuals had to navigate through a maze to hide in a shelter after encountering a predator. After displaying flight responses using the shortest route to the shelter, we tested each individual’s ability to find a new route to the shelter when the shortest route was blocked as a measure for behavioural flexibility. Furthermore, as a measure for their perceived fear after being exposed to a predator we recorded whether fish showed a strong fear response and hid inside the shelter or a weak fear response without entering the shelter.

Our study species *N. pulcher* is a highly social cichlid and in nature individuals live in groups consisting of dominant and subordinate individuals that are structured in a size-based, linear hierarchy (Taborsky, 2016; Wong and Balshine, 2011). Groups defend territories containing crevices and holes which are used as shelters for breeding and hiding from predators. Sexually mature *N. pulcher* feed on zooplankton in the water column which requires to stay far away from their shelters (Taborsky, 2016). Thus, flexibility in the way how individuals return to their shelter in case of danger is of utmost importance for their survival. Organizational effects of stress in early life are well researched in *N. pulcher* (Antunes et al., 2021a; Mileva et al., 2009) and pharmacologically blocked GRs during early life clearly altered the development of social competence and cognitive abilities (Nyman et al., 2018; Reyes-Contreras et al., 2019; Reyes-Contreras and Taborsky, 2022). Nevertheless, it is not known whether a short-term application of a GR antagonist immediately before a cognitive challenge mediates cognitive abilities and in particular behavioural flexibility.

Based on the role that GRs play in the negative feedback mechanism to terminate stress responses we formulated two predictions: (1) GR antagonist-treated fish show impaired behavioural flexibility, particularly after repeated exposures to a predator. (2) Effects that are mediated by GRs are specific to behavioural flexibility and the ability to cope with stress in terms of fear related behaviours and do not affect other behaviours such as the motivation of the fish to solve the task.

## Methods

### Subjects and housing

The experiment was conducted at the Konrad Lorenz Institute of Ethology, University of Veterinary Medicine, Vienna, Austria, under the licence (ETK-101/06/2019). All experimental tanks (45L, 30 x 50 x 30 cm) were equipped with a 2 cm layer of sand, a biological filter, a heater, and one-half clay flowerpot served as a shelter. The water temperature was kept at 27 ± 1°C with a light-dark regime of 13:11 h and a dimmed-light phase of 10 min. During the experiment all fish were fed commercial flake food 6 days a week.

Our study species was the cooperatively breeding cichlid *N. pulcher*, which is a model to study the evolution of vertebrate sociality highlighting their social and non-social behavioural flexibility (Taborsky, 2016; Wong and Balshine, 2011). These fish are endemic to Lake Tanganyika and groups always consist of a dominant breeding pair and related or unrelated helpers (Balshine et al., 2001; Heg et al., 2005; Taborsky, 1984, 1985). Helpers assist the breeding pair in raising the offspring by cleaning and fanning the eggs, cleaning the breeding shelters, and defending the common territory (Grantner and Taborsky, 1998; Taborsky, 1984, 1985). In larger territories each group member occupies and defends an own shelter. As an ecologically relevant stress stimulus, we used the piscivorous cichlid *Lepidiolamprologus elongatus* which occurs in sympatry with *N. pulcher* (Ochi and Yanagisawa, 1998). *L. elongatus* preys on all sizes of *N. pulcher*, except eggs, and is one of its most important predators (Groenewoud et al., 2016). Previous research indicates that *N. pulcher* can innately recognise this predator based on visual or olfactory cues (Fischer et al., 2014; Zöttl et al., 2013). In nature, groups of *N. pulcher* face a high predation pressure and individuals either attack or hide when predators intrude their territories, particularly when faced with a large predator such as *L. elongatus*.

Our test fish were laboratory-reared first- and second-generation offspring of wild-caught fish. We randomly selected 35 male and 35 female adult fish (N=70, mean standard length [SL; from the tip of the snout to the end of the caudal peduncle]: 7.2 cm, [range: 5.6-8.2 cm]) from the laboratory stock, where they had been kept before as groups separated by sex. The experiment was conducted in seven blocks over seven weeks with 10 test fish per block. Test fish were permanently marked using subcutaneous injections of Visible Implant Elastomer tags (VIE; Northwest 191 Marine Technology). Only fish which had been marked at least three days before the behavioural tests were included in the analyses reducing the final sample size to N=60.

We also randomly chose adult *L. elongatus* (N = 10, mean SL: 12.68 cm [range: 10.3-14.7 cm]) from the laboratory stock population, where they had been housed in pairs or small groups. We used five individuals for the first three blocks and another five individuals for the last four blocks. This ensured that predators behaved naturally throughout the experiment and were not exposed to excessive stress due to being caught and transported to the presentation tanks repeatedly (see below). During the experiment, each predator was kept in a separate 45L tank. These tanks were placed close to the tanks of the test fish but with a visual and olfactory separation. Body sizes of predators were obtained after the last block of the experiment was completed. All fish were kept under the same housing conditions and food regimes before and during the experiment.

### Experimental set-up

During the trials, we visually separated the experimental tanks using opaque dividers. The mazes used to carry out the detour trials remained in the experimental tanks throughout all the experimental phases (see below). The tanks were separated from the observer and the rest of the room using a dark curtain to minimize any disturbances from the surrounding area. All trials were recorded from above and from the front using two web cameras (Creative Live Cam Sync HD webcam) placed inside the curtain and connected to a computer outside of the curtain. Prior to each trial, a randomly chosen predator was transported to a small presentation tank (25 x 30 cm) filled with water and placed on a small movable rack. The front of the presentation tank – the part facing the experimental tanks – was covered with an opaque divider which could be moved with a string by the observer from outside the curtain. The divider was lifted before each trial for 60s to visually expose the test fish to the predator.

For all the trials, we used a transparent, U-shaped maze that prevented fish from using the maze as a shelter. The walls of the maze were covered with a mesh-like patterning that was fine enough to clearly see though. Halved clay flowerpots were used as shelters. The shelter could be reached using two routes: either via the direct route where fish only entered the arm of the maze that contains the shelter (‘shelter arm’, see Figure 1), or via a detour route where fish entered first the arm furthest away from the shelter (‘non-shelter arm’, see Figure 1) and had to cross to the other side in the back of the maze to enter the shelter arm. The middle arm of the maze was a dead end (Figure 1). The area in front of the maze (30 x 10 cm) that faced towards the predator was used as a starting area and could be separated from the rest of the tank with a removable opaque divider. Again, using a string the observer could remotely move the divider with minimal disturbance to the fish from outside the curtain. Biological filters and heaters were removed during the trials to ensure fish could only seek shelter inside the flowerpot.

**Figure 1:**
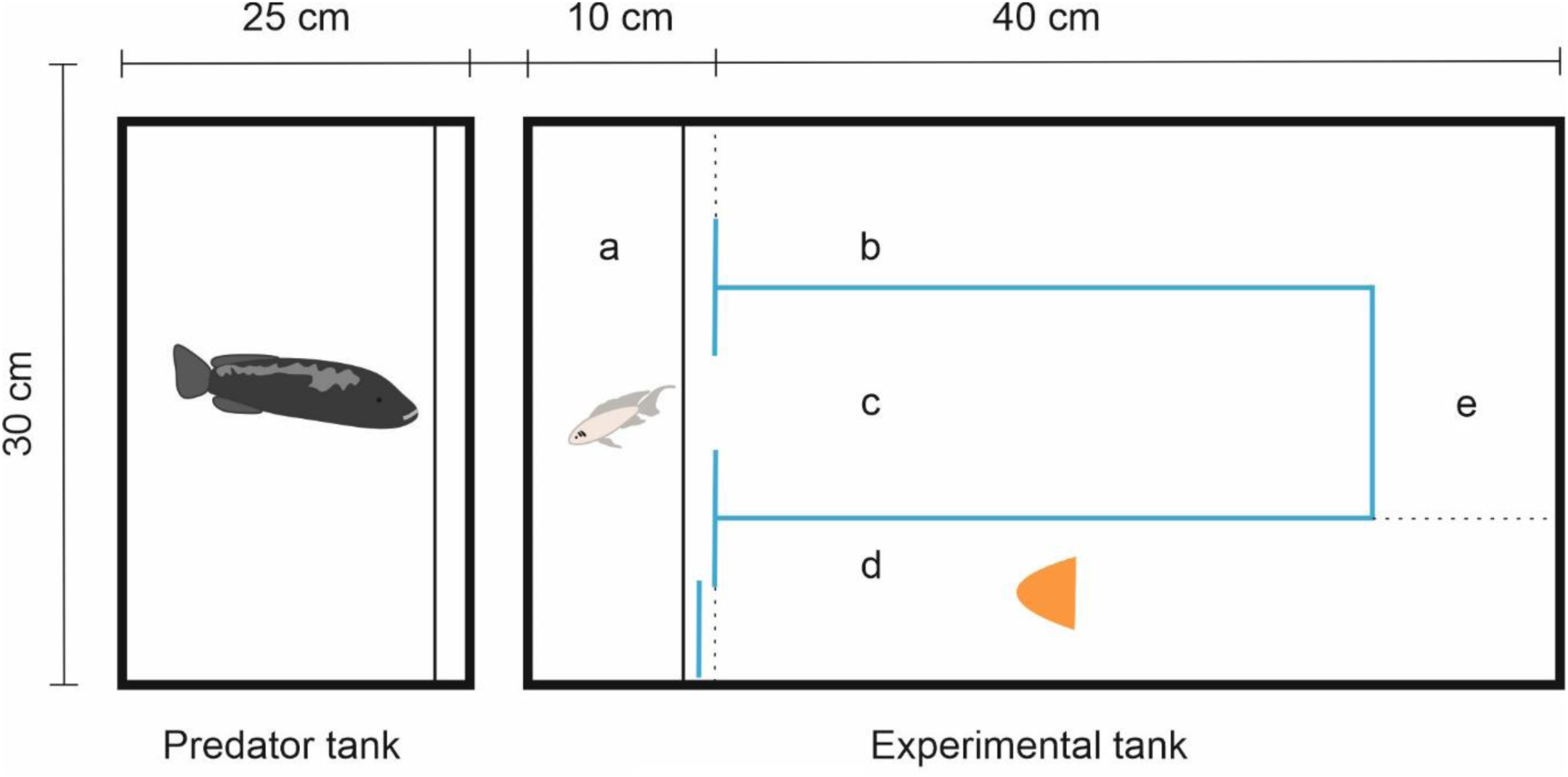
Experimental set-up consisting of one predator tank and a 45L experimental tank. The predator tank contained a removable opaque divider that could be remotely lifted during the predator presentations. The experimental tank was divided into five areas that are indicated with letters (a)-(e): (a) The start area which could be separated by a removable opaque divider from the rest of the experimental tank. (b) The non-shelter arm which served as the correct entrance in the detour trials. (c) The middle arm leading to a dead end. (d) The shelter arm that always contained a half clay flowerpot with the opening either facing towards the start area during the direct trials or with the opening facing away from the start area during the detour trials (as depicted here). (e) The back side arm which test fish had to use to enter the shelter arm in the detour trials. The U-shaped maze consisted of transparent Perspex material. A mesh was glued on top to make the maze clearly recognizable. Dashed lines indicate virtual lines that were used to record latencies and failed attempts from the video recordings (see Methods section for more details). The shortest route towards the shelter was either directly accessible (direct trials) or blocked (detour trials) with a barrier made from the same material as the maze.

### Experimental phases

Each block consisted of five distinctive experimental phases: a habituation phase (day 1, day 2), a pre-treatment direct phase (day 3), a treatment immersion phase (day 4, day 5), a post-treatment direct phase (day 6), and a detour phase (day 6).

#### Habituation

After fish were measured (SL) to the closest mm, each fish was individually transported into one of the experimental tanks. Each tank included a maze, and a shelter was placed in the shelter arm. Fish were allowed to habituate to the new environment for 48h. During the habituation phase we removed the opaque dividers between the experimental tanks to allow fish to visually interact with their neighbours.

#### Pre-treatment direct

After the habituation phase (day 3, 10a.m.), we tested whether the fish used the shortest route to the shelter after being exposed to a brief predator presentation. First, the observer gently guided the fish to the starting area using a small handheld net. Second, the observer lowered the divider to separate the start area from the rest of the tank. Third, the observer added an olfactory predator cue (15 ml of water from a predator tank) to the starting area and lifted the divider inside the predator tank to visually expose the fish to the predator for 60s. Fourth, directly afterwards, the observer lifted the divider separating the start area from the rest of the tank, and the fish could enter the maze and seek shelter. In this phase, the entrance of the shelter faced towards the starting area. Fish successfully completed this phase if they used the direct route to reach the shelter within 60s without any detours. Fish that did not use the direct route (N=8) were excluded from the experiment.

#### Immersion treatment

The subjects that successfully completed the previous phase (N=52), underwent a 48h period of either a mifepristone (N=26, females=14) or a control immersion treatment (N=26, females=13) starting on the next day (day 4, 8a.m.). The duration of the immersion treatment is well within the half-life of mifepristone (84h, see Molitch, 2022) and was chosen as a compromise between a maximised exposure time and a limited confinement of the fish in a small space. Half of the females and half of the males were randomly assigned to one of the two treatments, in a way that ensured that the observer was blind to the treatments of the fish within each block. For this a second person (SF) applied the treatments to the experimental tanks without the observer (ZF) present. The treatments were applied in the form of water baths following Nyman et al. (2018). In short, we dissolved mifepristone (RU 484) in dimethylsulfoxide at 50 mg ml-1, then serially diluted it in 0.1 M acetic acid (1:10), phosphate-buffered saline (1:100), and finally, diluted it in distilled water for an immersion concentration of 400 ng l^-1^. Controls were prepared with diluents without mifepristone. Both mifepristone and control solutions were prepared and stored in a −20C freezer one week prior to the experiment. The fish were immersed in 2L of water inside 3L glass containers that were placed inside the experimental tank.

#### Post-treatment direct

After the immersion treatments (day 6, 9a.m.) we repeated the previous direct phase to test whether (i) the fish memorised the shortest route to the shelter, and (ii) whether the immersion treatments influenced the motivation or memory of fish. Seven out of 52 fish did not take the direct route in this phase. This time all fish proceeded to the detour phase because they already used the direct route in the pre-treatment direct phase. This ensured that (i) all fish that proceeded to the detour phase had used at least once the direct route and (ii) the immersion treatments had no effect on the motivation of the fish to seek shelter.

#### Detour

For the detour trials (day 6, 10:30a.m.) we blocked the direct route to the shelter arm (Fig. 1) with a rectangular barrier (5 x 25 cm), made from the same material as the maze. The shelter entrance was turned to face away from the starting area. This ensured that fish approached the shelter from the same side irrespective of whether they entered the shelter arm via the direct or via the longer route (Figure 1). We used the same procedures as described for the pre-treatment direct phase. Each fish experienced 4 detour trials with a 2h recovery period between each trial.

### Video analysis

Video recordings were analysed using BORIS (Friard and Gamba, 2016). One trial (block 3, detour 1) was excluded from the analysis due to problems with the video quality. For each trial we recorded (1) the type of the test (post-treatment direct/detour 1/detour 2/detour 3/detour 4), (2) the latency to cross the shelter arm line, (3) the latency to enter the shelter, and (4) the strength of the fear response in terms of whether the fish entered the shelter or not. In the post-treatment direct trials, we additionally (5) recorded whether the fish used the shortest route to the shelter or not. To measure behavioural flexibility, we (6) recorded the number of failed attempts during the detour trials defined as crossing the middle arm line, attempting, and trying to cross the blocked shelter arm line, or crossing the non-shelter arm line but then returning to the starting area without entering the shelter arm. These variables allowed us to analyse the role of GRs in mediating motivation (variables 2 and 3), fear related behaviours (variable 4) and behavioural flexibility (variable 6). A fish was classified as having crossed into a new arm when both pectoral fins crossed the virtual line separating the two arms. We started measuring the latencies immediately after the predator exposure – i.e. immediately after the divider in the presentation tank was closed and the divider in the experimental tank was lifted. If fish did not cross any arm line or entered the shelter before the end of the experiment, we assigned maximum latencies (600s) in the respective trial.

### Statistical analysis

Data were analysed using R 4.1.1. (R Core Team, 2021) with the packages “lme4” (Bates et al., 2015) and “afex” (Singmann et al., 2021). We analysed treatment effects using Linear Models (LMs), Generalized Linear Models (GLMs), Linear Mixed Models (LMMs) and Generalized Linear Mixed Models (GLMMs). Residuals and Q/Q-plots of all LMs and LMMs were visually inspected, and the distributions of residuals were compared to a normal distribution using Kolmogorov-Smirnov and Shapiro tests. If residuals were not normally distributed, a log transformation was applied, and residuals again checked. All GLMs and GLMMs were checked for over-dispersion, but none required a correction. To obtain p-values, we used either the mixed() function in the package ‘afex’ with a Satterthwaite’s approximation for degrees of freedom, or the drop1() function to perform either a type II anova (for LM and LMM) or a χ^2^-test (for GLM and GLMM). Interactions, and control variables (sex and body size of test fish) were stepwise removed if non-significant (Engqvist, 2005) and reported p-values refer to the final model without non-significant interactions, and control variables. An overview of all full models can be found in Supplementary Table S1.

Except for the analyses of the probability to enter the shelter with increasing numbers of predator exposures (see below) we always included ‘Treatment’ (GR antagonist or control treatment), the body size and sex of test fish in the models. When analysing the detour trials, we additionally included test fish identity as a random effect to control for multiple observations on the same individual and the number of the detour trial as a fixed effect. If the interaction ‘No. of detour trial x Treatment’ was significant we compared the marginal means of both treatments within each detour trial separately (four comparisons) with a Tukey correction for multiple testing. To analyse the probability to enter the shelter with increasing numbers of predator presentations we used all trials that were done after the fish received their respective treatments (post-treatment direct, detour 1, detour 2, detour 3, detour 4). Fish identity was included as a random effect and the number of daily predator presentations as a continuous covariate. Initially we constructed two models, one that included a linear interaction between the number of daily predator presentations and treatment and one that included a non-linear, polynomial relationship with a degree of 2. We compared both models using a χ^2^-goodness of fit test. To foster comparability, we kept the structure of the models similar and did not include the body size or sex of fish in this analysis. To further investigate the significant non-linear interaction, we constructed two additional models separately for GR blocked and control fish. Both models included the identity of the fish as a random factor and the number of daily predator presentations as a linear predictor or as a non-linear polynomial predictor with a degree of 2.

## Results

### Does the application of a GR antagonist influence memory or motivation to seek shelter in the post-treatment direct trials?

We found no indication that the application of a GR antagonist influence memory or motivation because the majority of fish (87%) used the shortest route to the shelter again in the post-treatment direct trials and this is significantly higher than a random expectation of 50% (binomial test, p<0.01). Furthermore, the immersion treatments did not influence the choice of fish to use the short or the longer route (control: 4 out of 26 fish took the longer route; mifepristone: 3 out of 26 fish took the longer route; chi-square-test, χ2=0.17, p=0.68) or any other behaviours in the post-treatment direct trials. There were no significant differences between the two immersion treatments in the latency to cross the shelter line (F_1,50_=0.08, p=0.78; see factor: ‘Treatment’ in Table 1a), and the latency to enter the shelter (F_1,50_=0.1, p=0.77; see factor ‘Treatment’ in Table 1b).

**Table 1:** Latencies recorded during the post-treatment direct trials. (a) The latency to enter the shelter arm, and (b) the latency to enter the shelter were not significantly different in glucocorticoid receptor antagonist treated or control treated fish. The dependent variables were log transformed to obtain normally distributed residuals. Estimates are shown on a log scale and as differences to the reference level ‘control’ for factor ‘Treatment’. Body size and sex of test fish (N=52) did not predict any of the latencies and were subsequently removed from the models. F-values and p-values were obtained using likelihood ratio tests comparing models with or without the factor of interest.

| Factors | Estimate ± SE | Num. D.F. | Den. D.F. | F-value | P-value |
| --- | --- | --- | --- | --- | --- |
| **(a) Latency to enter the shelter arm** |  |  |  |  |  |
| Intercept | 2.92 ± 0.31 | - | - | - | - |
| Treatment | 0.12 ± 0.44 | 1 | 50 | 0.08 | 0.78 |
| **(b) Latency to enter the shelter** |  |  |  |  |  |
| Intercept | 3.29 ± 0.3 | - | - | - | - |
| Treatment | 0.12 ± 0.42 | 1 | 50 | 0.1 | 0.77 |

### Does the application of a GR antagonist influence behavioural flexibility in the detour trials during repeated exposures to a stressor?

GR blocked fish showed less behavioural flexibility during the detour trials than control fish. That is, the application of a GR antagonist influenced the number of errors that fish made over time with repeated exposures to predators (χ^2^=23.96, p<0.01, N=52; see interaction ‘No. of detour trial x Treatment’ in Table 2, Figure 2). Post-hoc analysis revealed that GR blocked fish made significantly more errors in the third detour trial compared to control fish (see Table 3 for pairwise comparisons). However, GR blocked fish did not show a longer latency to enter the shelter arm (F_1,50.11_=0.0, p=0.97, see factor ‘Treatment’ in Table 4a) or a longer latency to enter the shelter (F_1,48.98_=0.42, p=0.52, see factor ‘Treatment’ in Table 4b) than the control fish. Overall, all fish reduced the latency to enter the shelter arm with increasing predator exposures (F_1,152.25_=3.78, p=0.01; see factor ‘No. of detour trial’ in Table 4a), whereas this was not the case for the latency to enter the shelter (F_1,152.18_=0.6, p=0.61; see factor ‘No. of detour trial’ in Table 4b). Irrespective of the treatments, we found a positive relationship between body size and the latency to enter the shelter (F_1,48.9_=5.1, p=0.03; see factor ‘Body size’ in Table 4b) indicating that larger fish took longer to enter the shelter.

**Figure 2:**
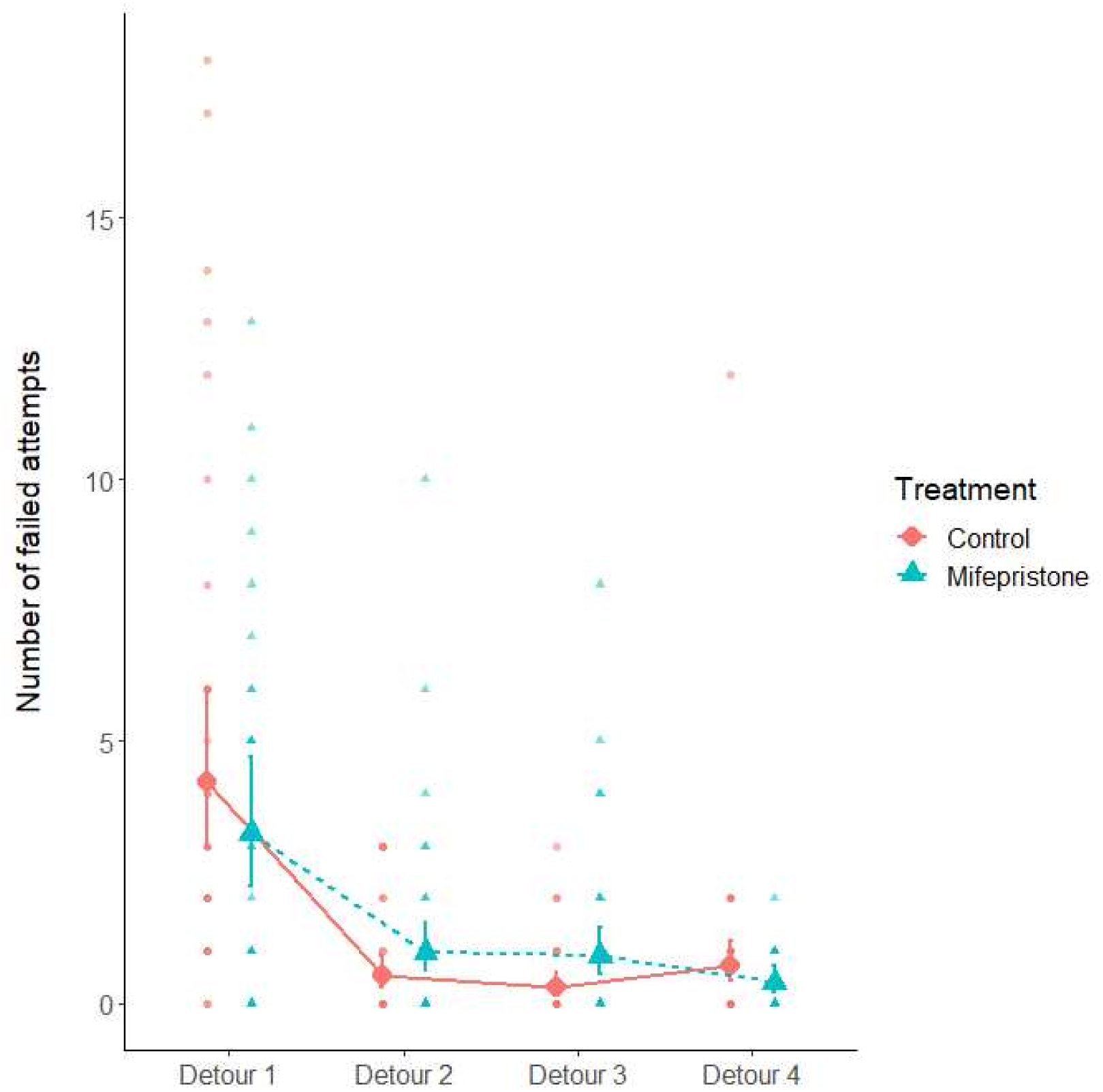
Behavioural flexibility as measured by the number of failed attempts in the detour trials. Glucocorticoid receptor (GR) antagonist treated fish showed a different pattern of failed attempts in the detour trials than control treated fish. GR blocked fish made more errors particularly during detour trial three. Individual data points (smaller circles and triangles in the background) and predicted values (larger circles and triangles in the front) are shown with 95% confidence intervals for control (red circles and solid line) and GR blocked (blue triangles and dashed line) fish.

**Table 2:** Number of failed attempts during the four detour trials. Fish treated with a glucocorticoid receptor (GR) antagonist showed a significantly different learning curve during the detour trials compared to control treated fish. (see significant interaction ‘No. of detour trials x Treatment’). Post-hoc tests revealed that GR blocked fish showed significantly more errors during detour trial 3 than control fish (see Table 3 for the pairwise comparisons). Estimates are shown on a log scale and as differences to the reference level ‘control’ for factor ‘Treatment’. Estimates for the factor ‘No. of detour trials’ and for the interaction ‘No. of detour trials x Treatment’ are presented as post-hoc comparisons in Table 3. Body size and sex of test fish did not predict the number of a failed attempts and were subsequently removed from the model. Χ^2^ and p-values were obtained from likelihood ratio tests comparing models with or without the factor of interest. N=52 (207 observations in total). P-values < 0.05 are highlighted in bold.

| Factors | Estimate ± SE | X<sup>2</sup>-value | P-value |
| --- | --- | --- | --- |
| Intercept | 1.44 ± 0.18 | - | - |
| Treatment | -0.26 ± 0.26 | 0.65 | 0.42 |
| No. of detour trial | - * | 288.56 | **<0.01** |
| No. of detour trial x Treatment | - * | 23.96 | **<0.01** |

\*Estimates are shown in Table 4

**Table 3:** Post-hoc comparisons with a Tukey correction of the significant interaction term in Table 2. Fish treated with a glucocorticoid receptor antagonist made significantly more errors during detour trail 3 leading to a significantly different error pattern in the detour trials (see Figure 2). Estimates are shown on a log scale, the direction of comparison within a contrast is left to right and the estimate values are shown as difference to the level at the left. N=52 (207 observations in total). P-values < 0.05 are highlighted in bold.

| Contrast | Estimate ± SE | z-value | P-value |
| --- | --- | --- | --- |
| **Detour 1** |  |  |  |
| Control vs Mifepristone | 0.26 ± 0.26 | 1 | 0.31 |
| **Detour 2** |  |  |  |
| Control vs. Mifepristone | -0.62 ± 0.37 | -1.7 | 0.09 |
| **Detour 3** |  |  |  |
| Control vs. Mifepristone | -1.15 ± 0.42 | -2.72 | **0.01** |
| **Detour 4** |  |  |  |
| Control vs. Mifepristone | 0.64 ± 0.41 | 1.57 | 0.12 |

**Table 4:** Latencies recorded during the detour trials. (a) The latency to enter the shelter arm, and (b) the latency to enter the shelter was not significantly different in glucocorticoid receptor antagonist treated or control treated fish. The dependent variables were log transformed to obtain normally distributed residuals. Estimates are shown on a log scale and as differences to the reference levels ‘control’ for factor ‘Treatment’ and ‘Detour 1’ for factor ‘No. of detour trial’. Sex of test fish did not predict the latencies and was removed from the models, whereas body size was only dropped in (a). F-values and p-values were obtained from likelihood ratio tests comparing models with or without the factor of interest. N=52 (207 observation in total). P-values < 0.05 are highlighted in bold.

| Factors | Estimate ± SE | Num. D.F. | Den. D.F. | F-value | P-value |
| --- | --- | --- | --- | --- | --- |
| **(a) Latency to enter the shelter arm** |  |  |  |  |  |
| Intercept | 4.92 ± 0.19 | - | - | - | - |
| Treatment | 0.01 ± 0.23 | 1 | 50.11 | 0.0 | 0.97 |
| No. of detour trial | - | 3 | 152.25 | 3.78 | **0.01** |
| Detour 2 | -0.49 ± 0.16 | - | - | - | - |
| Detour 3 | -0.39 ± 0.16 | - | - | - | - |
| Detour 4 | -0.36 ± 0.16 | - | - | - | - |
| **(b) Latency to enter the shelter** |  |  |  |  |  |
| Intercept | 1.71 ± 1.6 | - | - | - | - |
| Treatment | -0.14 ± 0.22 | 1 | 48.98 | 0.42 | 0.52 |
| Body size | 0.5 ± 0.22 | 3 | 48.9 | 5.1 | **0.03** |
| No. of detour trial | - |  | 152.18 | 0.6 | 0.61 |
| Detour 2 | -0.2 ± 0.2 | - | - | - | - |
| Detour 3 | -0.19 ± 0.17 | - | - | - | - |
| Detour 4 | -0.18 ± 0.17 | - | - | - | - |

### Does the application of a GR antagonist influence fear related behaviour with repeated exposures to a stressor?

When analysing the probability of fish to enter the shelter with increasing numbers of daily predator exposures, we found that a model containing a non-linear interaction between the number of daily predator exposures and treatment explained the data significantly better than a linear interaction term (χ2=15.37, p<0.01). GR blocked fish showed a linear reduction in the probability to enter the shelter with increasing numbers of daily predator exposures (z=-2.17, p=0.03, N=62; see factor ‘No. of predator exposure’ in Table 5a). In contrast, control fish showed a non-linear, U-shaped relationship between the probability to enter the shelter and the number of daily predator exposures [z=3.26, p<0.01, N=26; see factor ‘poly (No. of predator exposure, 2)’ in Table 5b]. Here, control fish entered the shelter more frequently during the initial and the last predator exposures than during any of the intermediate predator exposures (Figure 3).

**Figure 3:**
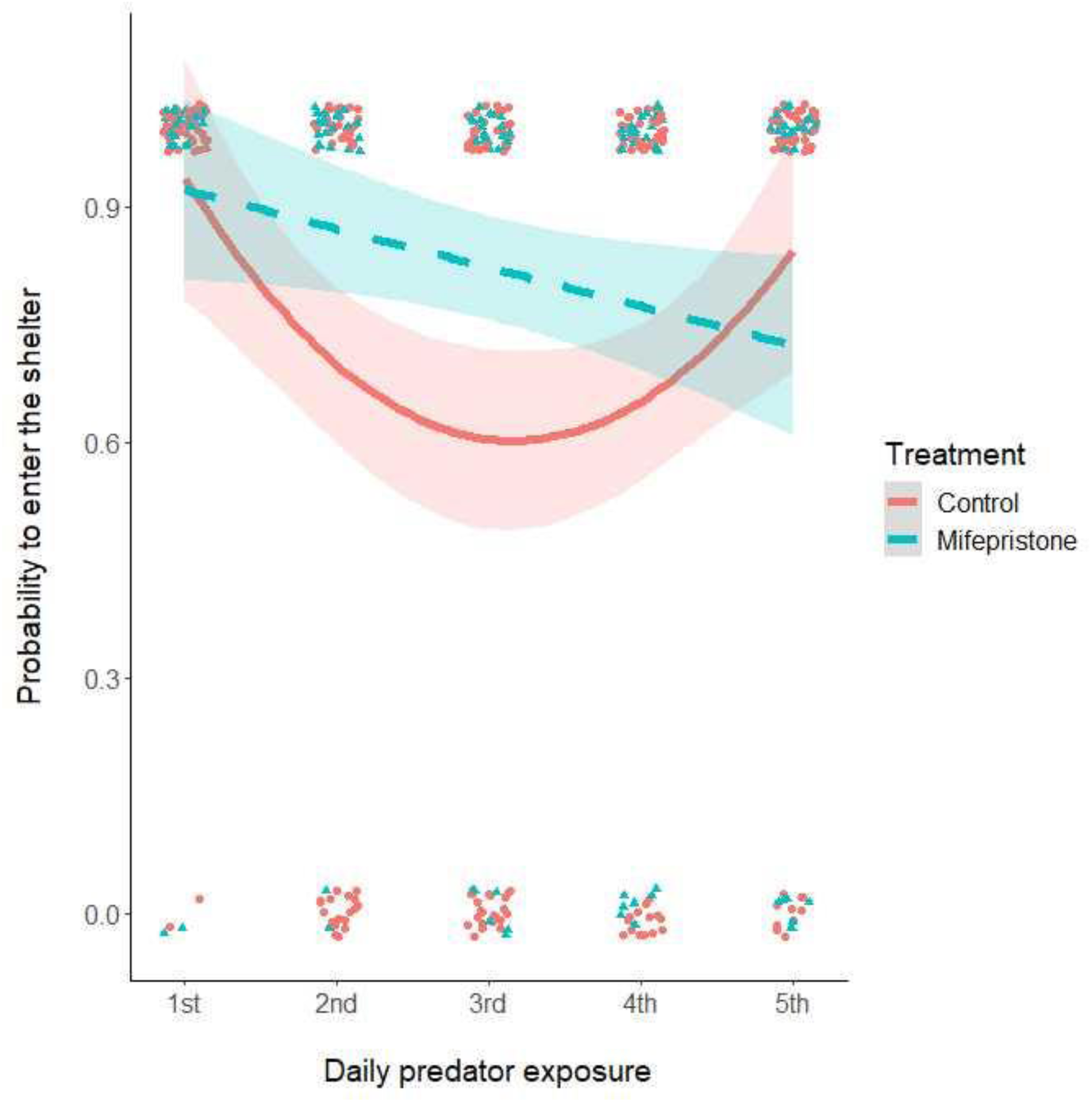
Fear-related behaviour measured by the probability to enter the shelter with increasing numbers of daily predator presentations for control treated (red circles and solid line) and glucocorticoid receptor (GR) antagonist treated (blue triangles and dashed line) fish. GR blocked fish showed a constant decrease in the probability to enter the shelter with successive predator presentations whereas control fish showed a nonlinear U-shape relationship. Circles and triangles represent raw data and lines represent model predictions with standard errors depicted as shaded areas.

**Table 5:**
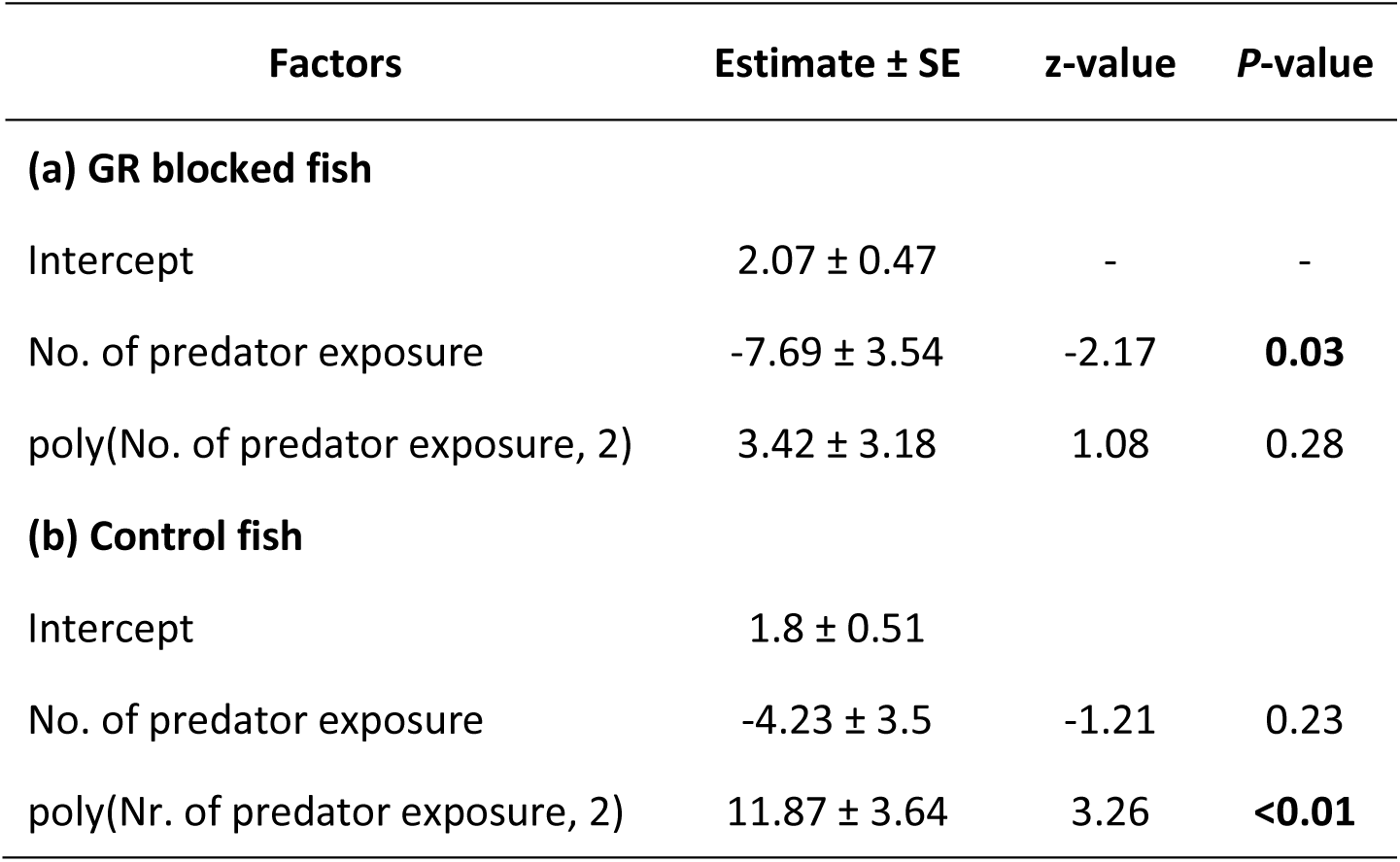
The probability to enter the shelter with increasing numbers of daily predator presentations. Fish treated with a glucocorticoid receptor (GR) antagonist showed a significantly different pattern over the course of the experiment than control treated test fish. Separate models were performed only after confirming that a non-linear interaction term between the number of daily predator presentations and treatment explained the data better than a linear relationship (see Results section). (a) GR blocked fish show a linear decrease in the probability to enter the shelter with increasing daily predator presentations, whereas (b) control fish show a significant U-shaped relationship (see also Figure 3). Estimates are shown on a logit scale. The factor ‘No. of predator exposure’ was included as a continuous variable either as a linear predictor or as a polynomial predictor (poly) of degree 2. N=26 (a: 129 observations; b: 130 observations in total).

| Factors | Estimate ± SE | z-value | P-value |
| --- | --- | --- | --- |
| **(a) GR blocked fish** |  |  |  |
| Intercept | 2.07 ± 0.47 | - | - |
| No. of predator exposure | -7.69 ± 3.54 | -2.17 | **0.03** |
| poly(No. of predator exposure, 2) | 3.42 ± 3.18 | 1.08 | 0.28 |
| **(b) Control fish** |  |  |  |
| Intercept | 1.8 ± 0.51 | | |
| No. of predator exposure | -4.23 ± 3.5 | -1.21 | 0.23 |
| poly(Nr. of predator exposure, 2) | 11.87 ± 3.64 | 3.26 | **<0.01** |

## Discussion

In the present experiment we applied a high dosage of a GR antagonist or a control immersion to groups of fish by a minimally invasive method, and subsequently tested their behavioural flexibility after repeated exposures to an ecologically relevant stressor. Fish treated with a GR antagonist (1) made more errors and (2) had a higher probability to enter the shelter during the intermediate compared to the initial and last predator exposures. Furthermore, the application of a GR antagonist did not influence the individual motivation to participate in the task or their spatial memory. More specifically, we did not find any behavioural differences during the post-treatment direct trials where test fish could enter the shelter using the shorter, direct route. We introduced this phase to control for potential influences of the application of a GR blocker on the motivation and memory of fish.

In our study fish had to find a new route to avoid a predator. Thus, our results show that GRs play an important role for behavioural flexibility in an ecologically relevant spatial task when repeatedly exposed to a mild naturalistic stressor. Behavioural differences between GR blocked and control fish were only apparent in the cognitively more demanding detour trials where inhibition control is required to successfully solve the task. This matches the results of Jungwirth et al. (2024), where the authors used a detour task in wild groups of *N. pulcher* and found that inhibition control was affected by the difficulty of the task. Although our study is not sufficient to reveal the precise underlying physiological mechanisms that drive the observed behavioural differences, the results highlight a potential role of fear in mediating behavioural flexibility: we found that the differences between GR blocked and control fish in terms of fear related behaviour (entering the shelter after predator exposure) and behavioural flexibility (number of failed attempts in the detour trials) followed the same pattern. In both behaviours, the difference between GR blocked and control fish was greatest during the second and third detour trial and both the number of failed attempts and the probability to enter the shelter were higher in GR blocked fish than in controls. GRs are distributed across the entire brain but GR activity in the basolateral amygdala and the ventral hippocampus have been specifically linked to the retention of fear memory (Donley et al., 2005).

Possibly, the difference in fear related behaviour was associated with increased concentrations of circulating GCs, in our case cortisol because previous studies showed that GCs are essential for fear related behaviours (for review see Schulkin et al., 2005b). Cortisol might have increased more in GR blocked fish due to their reduced capacity to terminate the stress response before encountering the next stressful predator exposure. However, another study in rainbow trout found a reduction in GC concentrations in GR blocked fish after exposure to a single stressor (Alderman et al., 2012). Nevertheless, it remains to be determined how GR activity influences GC concentrations after repeated exposures to the same stressor.

Although we did not directly measure cortisol, we observed a clear difference in fear related behaviours between the GR blocked and control fish. The response of control fish followed a U-shaped function with more fear related behaviour during the first and last predator presentations than during the intermediate presentations. The response of GR blocked fish followed a linear function with a continuous reduction of fear related behaviours from the first to the last predator presentation. We can think of three non-mutually exclusive explanations for this result. (1) In control fish the initial faster decline in fear related behaviour might reflect a fast habituation to the exposure to the stressor. In contrast, if GR blocked fish had increased cortisol levels due to the disruption of the negative feedback mechanism, they might have experienced chronic stress earlier which resulted in a slower habituation to the repeated stress exposure (Cyr and Romero, 2009). (2) Alternatively, the faster decline might indicate that control fish were better able to cope with the repeated exposure to stressors. A faster stress recovery may alter the potential responsiveness to a future stressor or aid in adapting to a chronic stressor (Sapolsky et al., 2000) (3) Finally, the subsequent increase of fear related behaviour in control fish might be explained by a potential ceiling effect for the facilitating action of stress during the acquisition of spatial information. Similar to our results, rats injected with an intermediate dosage of GCs performed better in a spatial learning task than rats injected with a low or a high dosage (Sandi and Pinelo-Nava, 2007). Further investigations are needed to disentangle the complex interplay between GCs, GRs and cognition particularly when repeatedly experiencing a stressor during development or later in life.

Mifepristone has several clinical applications and at low dosages it mainly acts as an antiprogestine whereas at higher dosages it mainly acts as a GR antagonist (Spitz, 2003). Our immersion treatment allowed us to apply a high dosage (400 ng l^-1^) of mifepristone which is more than fivefold than the highest dosage used in another study to investigate the effects of mifepristone as an antiprogestin in fish (Blüthgen et al., 2013). Thus, in our experiment, mifepristone acted mainly as a GR antagonist rather than as an antiprogestine, which limits the possibility that our results are explained by potential interactions between mifepristone and PRs. In line with this conclusion, we did not detect any sex specific effects on the observed behaviours. Nevertheless, progesterone is known to determine cognitive functions, particularly in humans (Griksiene et al., 2022) and future studies investigating the potential dose-dependent effects of mifepristone on cognitive abilities in *N. pulcher* are needed.

As in many animals, in *N. pulcher*, social and ecological experiences during early life are of particular relevance for the development of social and anti-predator behaviours, as well as for non-social behavioural flexibility (Bannier et al., 2017; Fischer et al., 2017). These organisational effects are most likely mediated by a life-long reprogramming of the physiological stress axis (Antunes et al., 2021a; Antunes et al., 2021b; Nyman et al., 2018) although an early-life manipulation of the stress axis using the same GR antagonist did not affect behavioural flexibility when adult individuals were tested in a reversal learning paradigm (Reyes-Contreras and Taborsky, 2022). In contrast, our study shows an activational effect on behavioural flexibility mediated by differences in the immediate stress response to a predator. Future studies focusing on both the organizational and the activational effects of the stress response on cognition, in particular on behavioural flexibility are needed to better understand the way how animals respond to naturalistic environmental challenges.

In conclusion, the current study provides novel insights into the mechanisms of adapting to environmental change. We used an ecologically relevant stressor and cognitive task to highlight the role of GRs in regulating fearful behaviour and mediating behavioural flexibility. We encourage further research to disentangle the interactions between stress responsiveness, cognitive abilities and associated behaviours in a natural context.

## Data availability

Data for this study are available at Figshare public repository: https://doi.org/10.6084/m9.figshare.24083952

## Acknowledgements

We thank Martina Krakhofer for help with animal husbandry, Roland Sasse for logistic support, Gopi Munimanda and Patricia Haubensak from the genetics laboratory for help with the immersion treatments and Stefan Graf for building the mazes. This study was financially supported by the Vienna Science and Technology Fund (CS18-042) and Z.F. was additionally supported by an Erasmus Mobility stipend.

## Author Contributions

Conceptualization & funding acquisition: K.H., B.T., L.F., S.T.; Data curation: S.F; Formal analysis: S.F., Investigation: S.F., Z.F.; Methodology: S.F., Z.F. S.T., Project administration: S.F., L.F., S.T.; Supervision: S.F., S.T.; Visualization: S.F.; Writing-original draft: S.F., Z.F.; Writing-review & editing: all authors (categories based on Contributor Role)

