## Supplementary Table S1 for "Does the stress axis mediate behavioural flexibility in a social cichlid, *Neolamprologus pulcher*?"

| Dependent variable | Factors | Covariates | Random factor | Transformation or Link | Interactions | Final model |
|---|---|---|---|---|---|---|
| Latency to enter the shelter arm (post-treatment direct) | Treatment<br>Sex of test fish | Body size | none | Log (LM) | none | Table 1a |
| Latency to enter the shelter (post-treatment direct) | Treatment<br>Sex of test fish | Body size | none | Log (LM) | none | Table 1b |
| Number of failed attempts (detour trials) | Treatment<br>No. of detour trial<br>Sex of test fish | Body size | Test fish ID | Log (GLMM) | Treatment x No. of detour trial | Table 2 |
| Latency to enter the shelter arm (detour trials) | Treatment<br>No. of detour trial<br>Sex of test fish | Body size | Test fish ID | Log (LMM) | Treatment x No. of detour trial | Table 4a |
| Latency to enter the shelter (detour trials) | Treatment<br>No. of detour trial<br>Sex of test fish | Body size | Test fish ID | Log (LMM) | Treatment x No. of detour trial | Table 4b |
| Whether a test fish entered the shelter (yes, no) (post-treatment direct and detour) | none | No. of predator exposure<br>poly(No. of predator exposure, 2) | Test fish ID | Logit (GLMM) | none | Table 6a, b |

**Table S1**: Information on the linear models (LM), linear mixed models (LMM), general linear models (GLM) and generalized linear mixed models (GLMM) analysed during this study, including the respective dependent variables, fixed factors, covariates and random factors, eventual data transformations performed to obtain normally distributed residuals, and any interaction terms included in the initial, full models. To obtain final models, interactions, the sex or the body size of test fish were removed if non-significant. Explanations of dependent variables: ‘Latency to enter the shelter arm’: the latency of fish to enter the shelter arm in the post-treatment direct or detour trials; ‘Latency to enter the shelter’: the latency of fish to enter the shelter in the post treatment direct or detour trials; ‘Number of failed attempts’: The number of failed attempts during the four detour trials; ‘Whether a test fish entered the shelter’: whether a fish entered the shelter after a predator exposure in the post-treatment direct and detour trials. Factor names: ‘Treatment’: whether fish were treated with a GR antagonist or a control solution (see Methods section for more details); ‘Sex of test fish’: The sex of fish; ‘No. of detour trial’: the number of detour trial (detour 1, detour 2, detour 3, detour 4). Covariate names: ‘Body size’: the body size of fish in cm; ‘No. of predator exposure’: how often a test fish was exposed to a predator presentation on the same day. Random factor names: ‘Test fish ID’: individual identification to correct for multiple observations on the same test fish.
